# Epigenetics of floral homeotic genes in relation to sexual dimorphism in the dioecious plant *Mercurialis annua*

**DOI:** 10.1101/481481

**Authors:** Janardan Khadka, Narendra Singh Yadav, Micha Guy, Gideon Grafi, Avi Golan-Goldhirsh

## Abstract

In plants, dioecy characterizes species carrying male and female flowers on separate plants and occurs in about 6% of angiosperms. To date, the molecular mechanism(s) underlying sexual dimorphism is essentially unknown. The ability of gender-reversal by hormone application suggests that epigenetics might play an important role in sexual dimorphism. Proteome analysis of nuclei derived from flower buds of females, males and feminized males of the dioecious plant *Mercurialis annua* revealed differentially expressed proteins related to nucleic acid binding proteins, hydrolases and transcription factors, including floral homeotic genes. Further analysis showed that class B genes are mainly expressed in male flowers, while class D genes, as well as *SUPERMAN*-like genes, were mainly expressed in female flowers. Cytokinin-induced feminization of male plants was associated with down-regulation of male-specific genes concomitantly with up-regulation of female-specific genes. No correlation could be found between the expression of class B and D genes and their DNA methylation or chromatin conformation. Thus, our results ruled out epigenetic control over floral identity genes as the major determinants regulating sexual dimorphisms. Rather, determination of sex in *M. annua* might be controlled upstream of floral identity genes by a gender-specific factor that affects hormonal homeostasis.

**Highlights:** Sex determination in *Mercurialis annua* is not related to epigenetics of floral homeotic genes but appears to be modulated by an unknown gender-specific regulator(s) that affects hormonal homeostasis.

## Introduction

The majority of angiosperms are hermaphrodites and monoecious (sexually monomorphic), whereby both male and female organs are found on the same individual plant. In contrast, only about 6% of the angiosperms are dioecious (sexually dimorphic) where male and female flowers are carried on separate individual plants (Renner and Ricklefs, 1995; Charlesworth, 2002). It has been hypothesized that dioecy has evolved independently from hermaphrodites through mutations in various families of plants (Charlesworth and Charlesworth, 1978; Charlesworth, 2002). Based on developmental aspects, dioecious plants are categorized into two types: type-I, unisexual flowers developed *via* abortion of reproductive organs that exhibit rudiments of the aborted organs, and type-II, to which *Mercurialis annua* belongs, where unisexual flowers do not have rudiments of the opposite sex (Mitchell and Diggle, 2005).

Most studies related to the regulation of flower development were performed in model plants such as *Arabidopsis thaliana* that has hermaphroditic flowers with four concentric whorls: sepals, petals, stamens and carpels. The homeotic genes that regulate the development of such flowers were described by the ABCDE model (Coen and Meyerowitz, 1991; Krizek and Fletcher, 2005). These gene classes were termed according to the whorl in which they function: class A genes *APETALA1* (*AP1*) and *APETALA2* (*AP2*), class B genes *APETALA3* (*AP3*), *TOMATO MADS-BOX GENE6* (*TM6*) and *PISTILLATA* (*PI*), class C gene *AGAMOUS* (*AG*), class D genes *SHATTERPROOF1* (*SHP1*), *SHATTERPROOF2* (*SHP2*) and *SEEDSTICK* (*STK*) also known as *AGAMOUS*-*like1* (*AGL1*), *AGAMOUS*-*like5* (*AGL5*) and *AGAMOUS*-*like11* (*AGL11*), respectively, and class E genes *SEPALATA1* (*SEP1*) to *SEPALATA* (*SEP4*). Most of these genes encode MIKC-type MADS-box domain containing transcription factors. The MADS-box proteins form homo- and/or heterodimers that bind specific DNA sequences, CC(A/T)_6_GG, called CArG box and the pairs of CArG-boxes are brought into close proximity by DNA looping (Nurrish and Treisman, 1995; Davies *et al*., 1996; Riechmann *et al*., 1996; Mendes *et al*., 2013). The protein dimers further form functionally active tetrameric protein complex termed, ‘floral quartet’, that specifically controls differentiation of flower whorls (Theissen and Saedler, 2001; Smaczniak *et al*., 2012). Accordingly, combination of A+E genes specify sepals, A+B+E genes specify petals, B+C+E genes specify stamens, C+E genes specify carpels and D+E genes specify ovule identity (Theissen and Saedler, 2001; Soltis *et al*., 2007).

As floral homeotic MADS-box genes control the expression of other regulatory genes, these genes are considered as key factors for development of the floral organs, both perianth parts and sex organs (Wuest *et al*., 2012; O’Maoileidigh *et al*., 2013; Stewart *et al*., 2016). However, the regulation of these genes is not fully understood. In *Arabidopsis*, *SUPERMAN* (*SUP*) is required for proper development of the reproductive organs inasmuch as mutation of the *SUP* gene has led to extra stamens formation at the expense of carpel development (Jacobsen and Meyerowitz, 1997; Sakai *et al*., 1995). The *SUP* transcription factor is proposed to act as a negative regulator of class B genes (i.e., *AP3* and *PI*) to maintain boundaries between the stamen and the carpel whorls (Bowman *et al*., 1992; Yun *et al*., 2002; Prunet *et al*., 2017) and its role appears to be conserved among dicot and monocot plants (Nandi *et al*., 2000). Additionally, *SUP* gene is required for development of the outer integument of the ovule (Gaiser *et al*., 1995). In dioecious *Silene latifolia*, *SUPERMAN* orthologous gene, *SlSUP* was associated with female flower development (Kazama *et al*., 2009). Ectopic expression of *SUP* in tobacco plants was shown to induce increased feminization via enhancing cytokinin related processes (Nibau *et al*., 2011). A recent report addressing gender-specific methylation in the sex determining region of *Populus balsamifera* identified the PbRR9 gene showing a clear pattern of gender-specific methylation. PbRR9 encode for a protein member of the two-component response regulator (type-A) gene family involved in cytokinin signaling (Brautigam *et al*., 2017).

Only limited research has been made to elucidate the role of floral homeotic genes in sexually dimorphic dioecious plants. This is surprising in view of the apparent advantage of separation of the reproductive organs between female and male plants, which makes it experimentally more amenable to investigation of the developmental regulation of each gender in plants. We have shown recently dimorphic responses of *M. annua* plant genders to stress that may be attributed to female plants’ capacity to survive stress and complete the reproductive life cycle (Orlofsky *et al*., 2016). A few of the more studied dioecious species include *Silene latifolia* and *Rumex acetosa* of the type-I flowers and *Thalictrum dioicum* and *Spinacia oleracea* of the type-II flowers. It was shown that in the case of *S*. *latifolia* and *R*. *acetosa*, class B and C floral organ identity genes are expressed early in development of male and female flowers (Hardenack *et al*., 1994; Ainsworth *et al*., 1995), while in *T. dioicum* and *S*. *oleracea*, class B and C genes are differentially expressed at floral initiation (Di stilio *et al*., 2005; Pfent *et al*., 2005). Sather *et. al*. (2010) showed that silencing of class B genes in *S. oleracea* is able to alter the floral gender of males into hermaphrodites or females due to transformation of stamens into carpels.

The annual dioecious (type-II) *M. annua* L. (Euphorbiaceae) is a unique model plant for the study of dioecism, since it has a short life cycle, which enables molecular-genetic studies, in contrast to most dioecious plants, which are woody perennials. It is a common roadside herb native to the drought and high-sunlight prone Mediterranean basin, which has spread into Europe, North America and Australia (Durand and Durand, 1991; Pannell *et al*., 2008). The diploid species (2n=16) is a strictly dioecious, while polyploid species are not (Thomas, 1958; Durand and Durand, 1991; Pannell *et al*., 2008). The dioecious *M. annua* has an interesting genetic system of sex determination lacking heteromorphic sex chromosome. Identification of male specific molecular markers and recent genetic analyses have revealed that males possess homomorphic XY chromosomes, but the molecular mechanism of sex determination is not clear yet (Khadka *et al*., 2002, 2005; Russell and Pannell, 2015; Veltsos *et al*., 2018). Furthermore, sex expression in *M. annua* can be reversed by exogenously applied plant growth hormones. Accordingly, auxins have a masculinizing effect while cytokinins have a feminizing effect (Delaigue *et al*., 1984). The ability of gender-reversal by hormone application suggests that the gene(s) required for the development of both type of flowers are genetically functional but might be restrained by epigenetic means in the floral primordia, even when lacking vestiges of the opposite sex, thus being still sexually bi-potent.

Here we attempted to study the epigenetic regulation of floral identity genes in *M. annua* and the relationship with sex determination. We report that differential expression of floral homeotic genes was associated with sexual dimorphism in *M. annua* and cytokinin was involved in their transcriptional control. The possible involvement of epigenetic regulation of the examined floral genes was ruled out.

## Materials and methods

### Plant growth condition

Dioecious *Mercurialis annua* (Euphorbiaceae), Belgian origin, was used in this study. Seeds were sown in trays containing standard gardening soil and the seedlings were transplanted into pots (2.5 L) and grown in a controlled climate growth chamber at 27 °C with photoperiod regime of 14 h light/10 h dark and light intensity of approximately 400 µmol m^-2^ sec^-1^.

### Feminization of male plants by 6-benzylaminopurine treatment

At the onset of flowering (about 25-day-old plants), male plants were separated from female plants. Feminization of the isolated male plants was done by spraying 1 mg L^-1^ 6-benzylaminopurine (BAP) three times daily as described (Durand and Durand 1991; Khadka *et al*., 2005). Inflorescence bud samples were collected and either used immediately for nuclei isolation or stored at −80°C until analyzed.

### Proteomic analysis

Nuclei isolated from flower buds were subjected to proteome analysis by the proteomic services of The Smoler Protein Research Center at the Technion, Israel. The samples were digested by trypsin, analyzed by LC-MS/MS on Q-Exactive (Thermo) and identified by Discoverer1.4 software against *Ricinus communis, Jatropha curcas* and *Arabidopsis* protein databases. All the identified peptides were filtered with high confidence, top rank and mass accuracy. High confidence peptides were passed the 1% FDR threshold (FDR =false discovery rate, is the estimated fraction of false positives in a list of peptides). The peak area on the chromatogram of the protein was calculated from the average of the peptides from each protein. PANTHER classification tool was used for categorization of differentially expressed proteins (Mi *et al*., 2013).

### Nucleic acid extraction and cDNA synthesis

Genomic DNA was extracted using the PureLink Genomic DNA Mini Kit according to the manufacturer’s protocol (ThermoFisher Scientific). Total RNA was extracted using the RNeasy Mini Kit (Qiagen). The first strand of cDNA was synthesized from 1 µg DNase (Epicentre)-treated total RNA using Verso cDNA Synthesis Kit (ThermoFisher Scientific).

### Isolation of genes and partial promoter sequences

Floral homeotic cDNA clones were prepared by PCR using *M. annua* flower cDNA as template and appropriate degenerate primers (based on conserved regions of *A. thaliana*, *Ricinus communis* and *Jatropha curcas*; for primer sequences see Supplementary file 1) for the recovery of class B (*AP3*, *PI*, *TM6*), class C/D (*AG*, *AGL5* and *AGL11*), as well as two *SUPERMAN*-like (*SUP*-like) gene products. PCR conditions were: 95 °C, 2 min; 40 cycles of 95 °C, 30 s; 65-45 °C, 30 s; 72 °C, 60 s; followed by 72 °C, 5 min. The PCR products were purified using QIAquick gel extraction kit, then cloned into pJET1.2 plasmid vector (ThermoFisher scientific) and sequenced at the Biotechnology Center, Ben-Gurion University of the Negev, Beer-Sheva, Israel.

To obtain full cDNA sequence, 3’-RACE was performed as described by Yadav *et al*. (2012) and 5’-RACE was performed using a 5’-Full RACE Core Set kit (TaKaRa). The purified PCR products were directly sequenced as above.

Based on phylogenetic analyses (see supplementary text for details), the class B genes were designated as *MaPI* for *PI* ortholog, *MaAP3* for *AP3* ortholog, *MaTM6* for *TOMATO MADS BOX GENE6* ortholog. The AGAMOUS-like genes were designated as *MaAG1* for *AGAMOUS* ortholog (class C), *MaAGL1* for *STK*/*AGL11* ortholog (class D) and *MaAGL3* for *SHP2*/*AGL5* ortholog (class D). NCBI GenBank accession numbers: KR781112-6.

The upstream promoters of *MaAP3*, *MaAGL1*, *MaPI*, *MaSL1* and *MaSL2* were isolated by semi-random sequence walking strategy modified from Aquino and Figueiredo (2004). Briefly, a gene specific primer was used for linear amplification of specific DNA segment for 20 high stringency cycles (95 °C, 30 s; 60 °C, 30 s; 72 °C, 2 min). Then random walking primer was added and a low stringency cycle (95 °C 30 s, 35 °C 30s, 72 °C 2 min) was used for unspecific binding and amplification. Then, 30 high stringency cycles were used for exponential amplification. The desired fragments were screened by semi-nested PCR using asymmetrical ratio (1:5) of walking primer and nested gene specific primer. The products of interest were purified, cloned and sequenced as above.

For reference, a 135 bp of Actin gene was amplified using primers designed from conserved region of mRNA of *J. curcas*, *R. communis* and *Populus trichocarpa*. The amplified product of *M. annua ACTIN* (*Act*) gene was confirmed by direct sequencing from both ends.

### Gene expression analysis

Quantification of the gene expression level was done by quantitative or semi-quantitative RT-PCR analysis using gene specific primers. qPCR was carried out using Perfecta SYBR green supermix (Quanta Biosciences). Amplification was conducted on an Applied Biosystems^®^ 7500 Real-Time PCR Systems. All reactions were performed from three biological samples and each with three technical replicates. The PCR conditions were: 94 °C for 15 s, 40 cycles of 94 °C for 5 s, 60 °C for 30 s. Each reaction was normalized against the expression of *Actin* gene. The relative changes in gene expression was calculated using the 2^-ΔΔ*C*T^ method (Livak and Schmittgen, 2001).

### Micrococcal nuclease accessibility assay

Micrococcal nuclease (MNase) accessibility assay was performed as described (Zhao *et al*., 2001). Nuclei prepared from male and feminized *M. annua* flower buds were incubated with MNase for various durations, and the DNA was resolved on agarose gel. MNase treatment resulted in a nucleosomal ladder. The recovery of DNA after MNase treatment was checked by PCR.

### DNA methylation analysis

For cytosine methylation analysis, chop-PCR (methylation-sensitive enzyme digestion followed by PCR) and bisulfite sequencing were performed as described (Yadav *et al*., 2018). In chop-PCR, genomic DNA was treated with methylation sensitive restriction enzymes *Hpa*II or *Msp*I and subjected to PCR to amplify various gene fragments containing the restriction site ‘CCGG’.

Bisulfite conversion was done by adding a mixture of sodium bisulfite, hydroquinone and urea, and incubated at 55 °C for 16 hrs. The samples were desalted using PCR purification kit and desulfonated by adding NaOH to a final concentration of 0.3 M. Then, DNA was purified by QIAquick PCR purification kit (Qiagen). The bisulfite converted DNA was used for PCR amplification of promoter and gene-body of MaAP3 and MaSL1 genes. The PCR products were cloned into pJET1.2 vector. At least 10 individual clones from each region were sequenced by Macrogen, Netherlands. The sequences were analyzed and scored using Kismeth online service (Gruntman *et al*. 2008).

### Statistical analysis

The data representing the average values of three biological replicates each with three technical chemical replicates and error bars representing the standard deviations were graphed. Student’s t-test was used to determine the statistical significance of differences at the p-level of 0.01. Error bars indicate SE of the mean (n=3).

## Results

### Feminization of male *Mercurialis annua*: setting up the experimental system

The female and male *M. annua* plants have distinct inflorescence morphology (Fig. 1A and 1B). In female plants, flowers developed directly at leaf axils with short pedicels, while in male plants, clusters of flowers developed on long pedunculated inflorescences. Feminization of male flowers by the cytokinin, 6-Benzylaminopurine (BAP), caused development of female flowers that yielded fertile seeds (Fig. 1C) on male inflorescences (Khadka *et al*., 2005).

**Figure 1.**
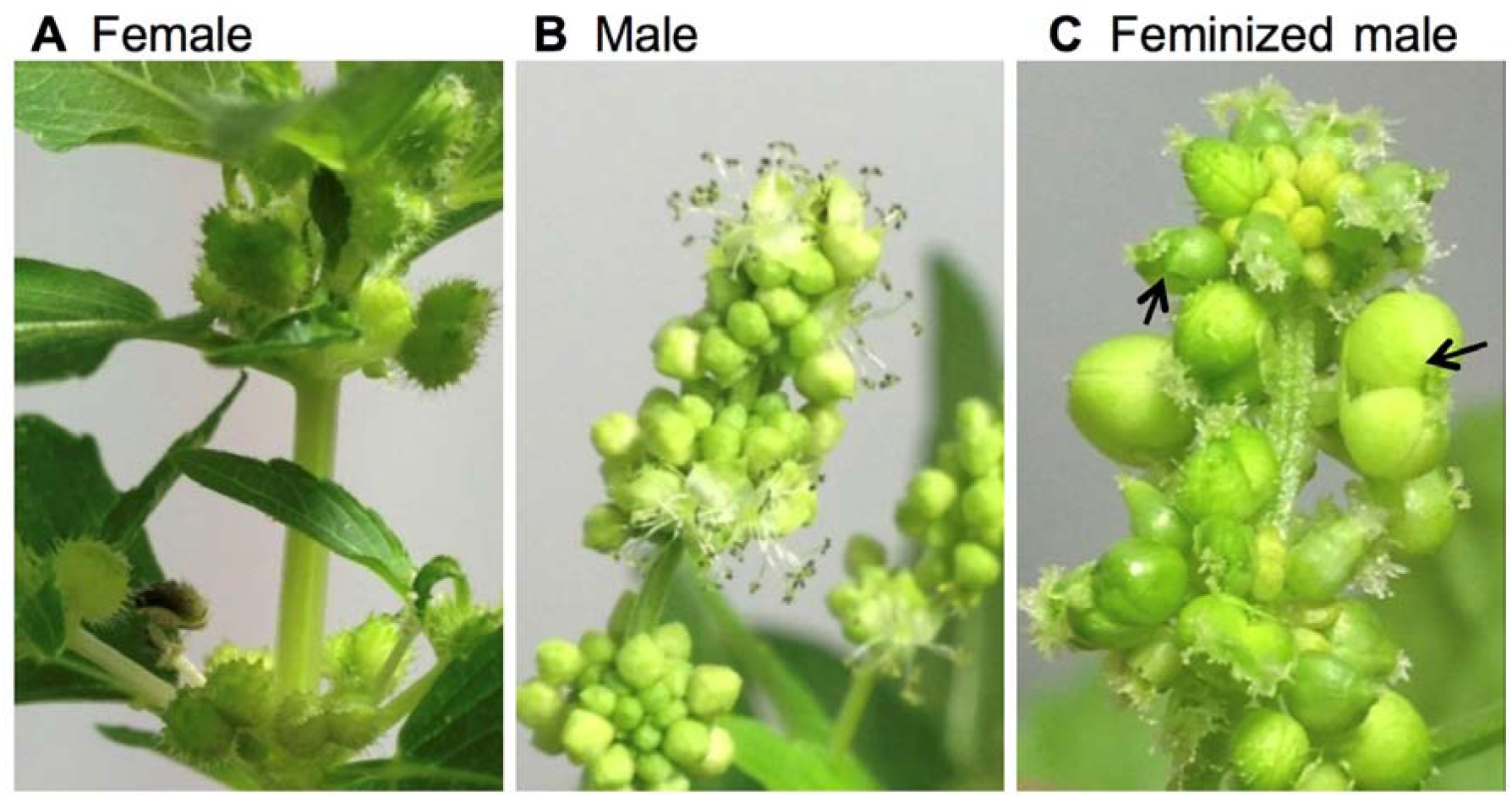
Morphological observation of dioecious *M. annua*. (**A**) Female plant. (**B**) Male inflorescence. (**C**) Feminized male inflorescence, BAP-hormone was sprayed 3 times daily for 4 weeks. Note that feminized male produced bi-carpellet flowers (some are indicated by arrows).

### Proteome analysis of flower bud nuclei

To identify regulatory genes involved in sexual dimorphism and BAP induced sex alteration of *M. annua*, we performed proteome analysis of nuclear proteins derived from flower buds of female, male and males treated with BAP for 4, 8, 12 and 16 days. The proteome data showed a total of 1443 proteins. Nuclear proteins including core histone proteins H2A, H2B, H3 and H4 displayed the highest intensities among the proteins identified in this analysis (Supplementary file 2, S1). The major difference between the genders was that 52 proteins that were present in female flowers were absent in male flowers, while 244 proteins that were present in males were absent in female flowers. Among the 52 proteins expressed only in female flowers, 49 proteins were up-regulated in feminized males (Supplementary file 2, S2). Out of the 244 male-specific proteins, 84 proteins were disappeared in the course of feminization (Supplementary file 2, S3). The change in protein expression was as follows: 39 proteins disappeared after 4 days of BAP treatment, 15 after 8 days, 12 after 12 days and 18 proteins disappeared after 16 days of BAP treatment.

Multiple classes of proteins were identified by categorization analysis of the differentially expressed proteins in feminized male. The major up-regulated protein classes were nucleic acid binding proteins, transcription factors and cytoskeleton proteins (Fig. 2A), and the major down-regulated protein classes were hydrolases, nucleic acid binding proteins, ligases and transferases (Fig. 2B). Among differentially expressed proteins, four floral organ identity MADS-box transcription factors were identified. The class E proteins, *SEPALLATA1* (*SEP1*) and *SEPALLATA3* (*SEP3*), and the class D protein *SHP2*/*AGL5* were up-regulated during feminization, reaching a maximum at day 16 (Fig. 2C, D and E). In contrast, the class B protein *PISTILLATA* (*PI*) was down-regulated within 4 days and disappeared completely afterwards (Fig. 2F).

**Figure 2.**
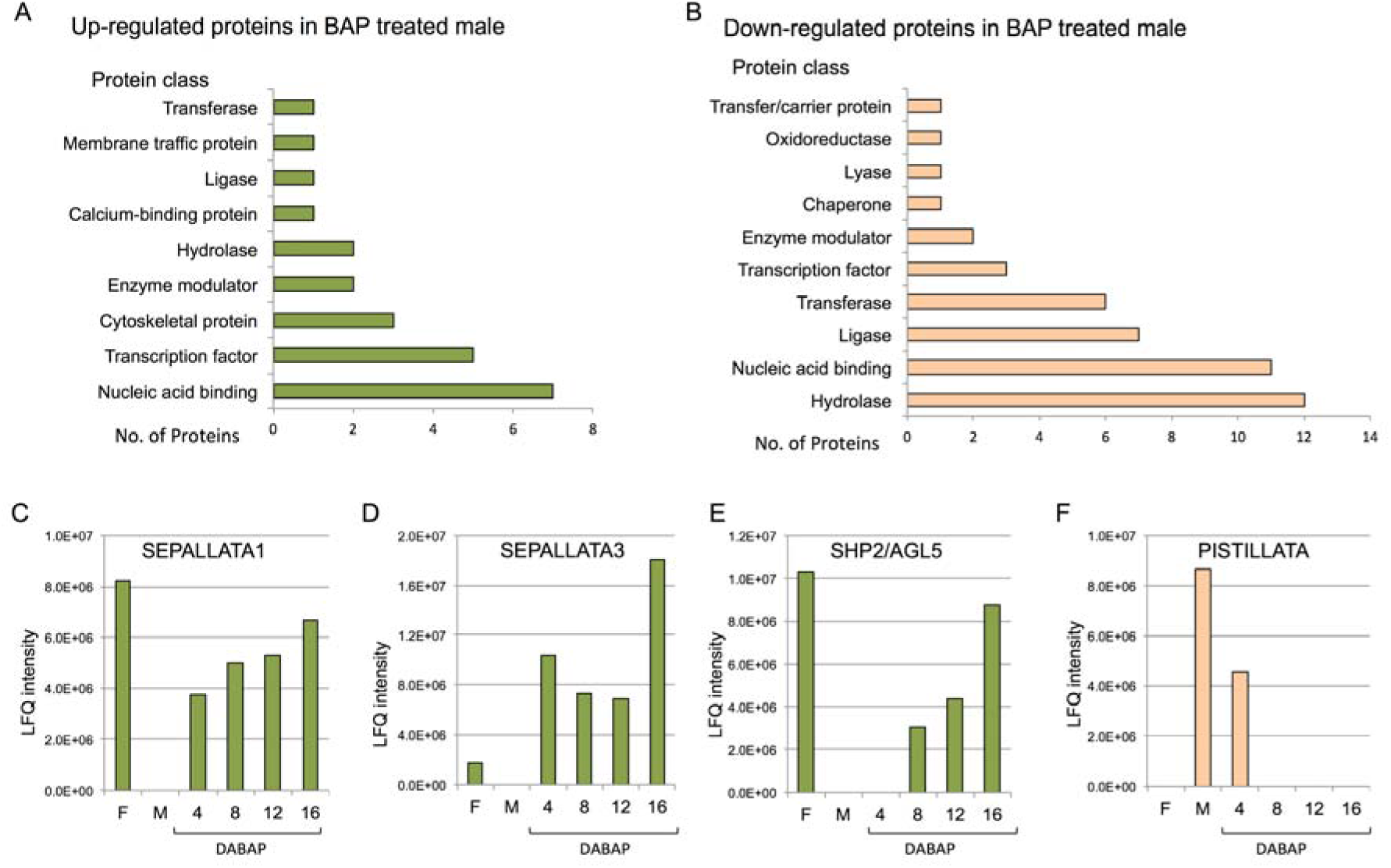
Proteome analysis of nuclei isolated from female, male and BAP treated male flower buds. Categorization analysis of down-regulated (**A**) and up-regulated (**B**) proteins following male feminization. The LFQ intensity of SEPALLATA1 (**C**), SEPALLATA3 (**D**), SHP2/AGAMOUS-like5 (**E**) and PISTILLATA (**F**) proteins in female, male and BAP treated males for 4, 8, 12 and 16 days, is shown. F, Female; M, Male; DABAP, Days after initiation of BAP treatment. Note that LFQ (label-free quantification) intensity reflects the relative amounts of the proteins, which was calculated using peptide intensities normalized between the samples.

### Differential expression of floral homeotic genes

The proteome data prompted cloning and analysis of *M. annua* orthologs of floral homeotic MADS-box genes. The RNA expression pattern of the isolated genes in female and male flowers, at bud and opened-flower stages was investigated (Fig.3). The class B genes *MaPI* and *MaAP3* were highly expressed in male flowers and poorly expressed in female flowers (Fig. 3A and 3B). The class C gene, *MaAG1* was strongly expressed with similar level in female and male flowers (Fig. 3C). In contrast, the class D genes *MaAGL1* and *MaAGL3* were strongly expressed in female flowers and poorly expressed in male flowers (Fig. 3D and 3E). Moreover, the expression level of most floral genes was significantly different (p<0.01) between the floral bud stage and the open flower stage; the expression of *MaAP3* and *MaAG1* was higher at flower bud developmental stage, while the expression of *MaPI* and *MaAGL3* was higher at open flower developmental stage (Fig. 3A, B, C and E).

**Figure 3.**
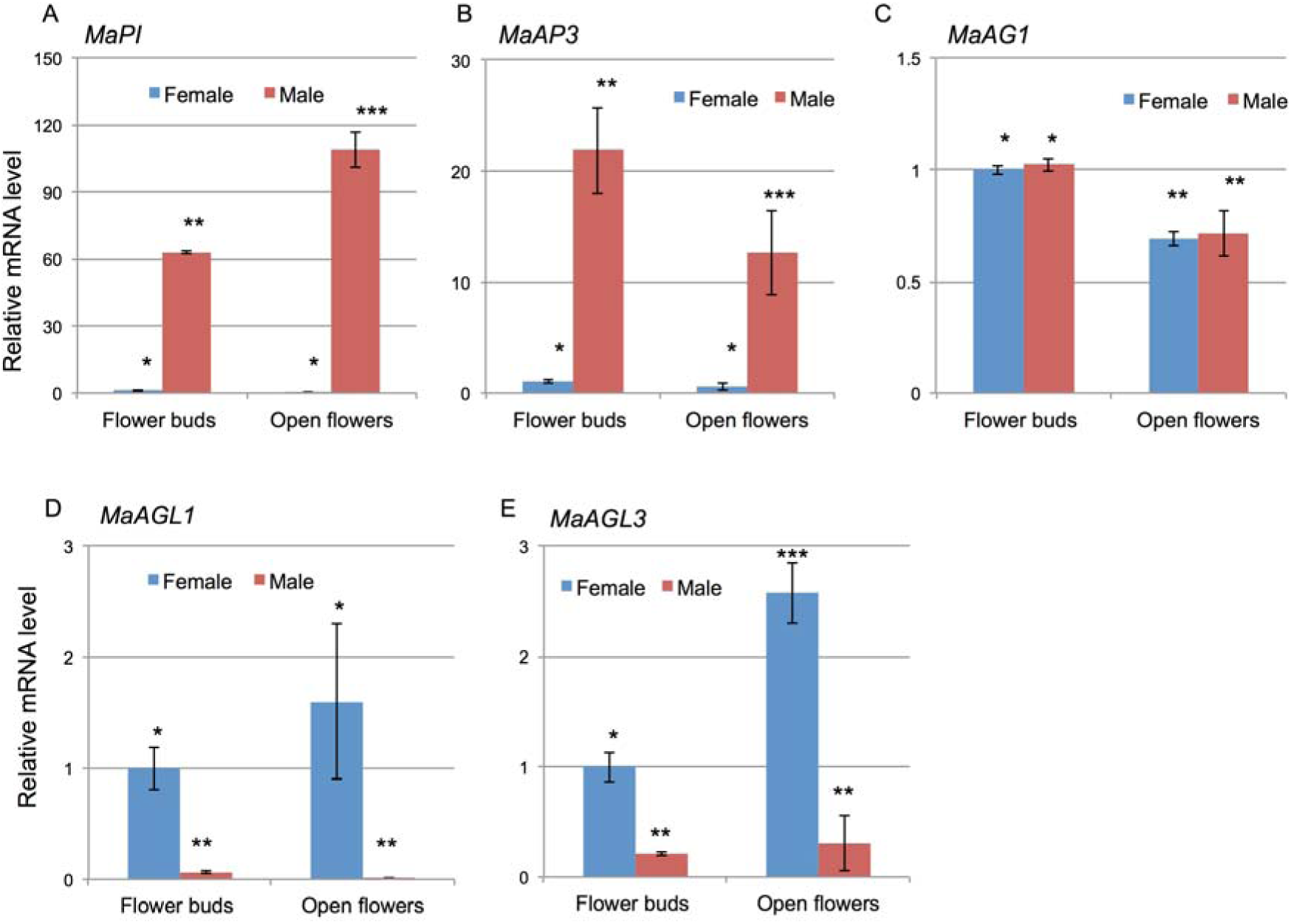
Expression of MADS-box genes in flower buds and open flowers of female and male plants of *M. annua*. Relative expression of (**A**) *MaPI*, (**B**) *MaAP3*, (**C**) *MaAG1*, (**D**) *MaAGL1* and (**E**) *MaAGL3* genes determined using RT-qPCR. Y-axis shows relative transcript level of genes normalization to *Actin* gene. The values are average of three biological replicates. Values denoted by different numbers of asterisks are significantly different (Students *t*-test, P < 0.01). Error bars indicate the standard error of the mean (n=3).

The flower organ specificity of gene expression (Fig. 4) showed that *MaPI* and *MaAP3* were almost exclusively expressed in male flowers, noteworthy that *MaPI* was also strongly expressed in peduncles. *MaTM6* gene expression was evident in flowers of female and male plants. The *MaTM6* expression was relatively higher in flowers and low in vegetative organs of female plants. In the male, *MaTM6* expression was highest in flowers, moderate in leaves and peduncles, and very low in stem and roots. *MaAGL1* and *MaSL1* were exclusively expressed in flowers of female plants. *MaAG1* was expressed at moderate level in flowers of both sexes, and at lower level in peduncle of male plants. *MaAGL3* was highly expressed in flowers of female and slightly lower expression in flowers and peduncles of male plants.

**Figure 4.**
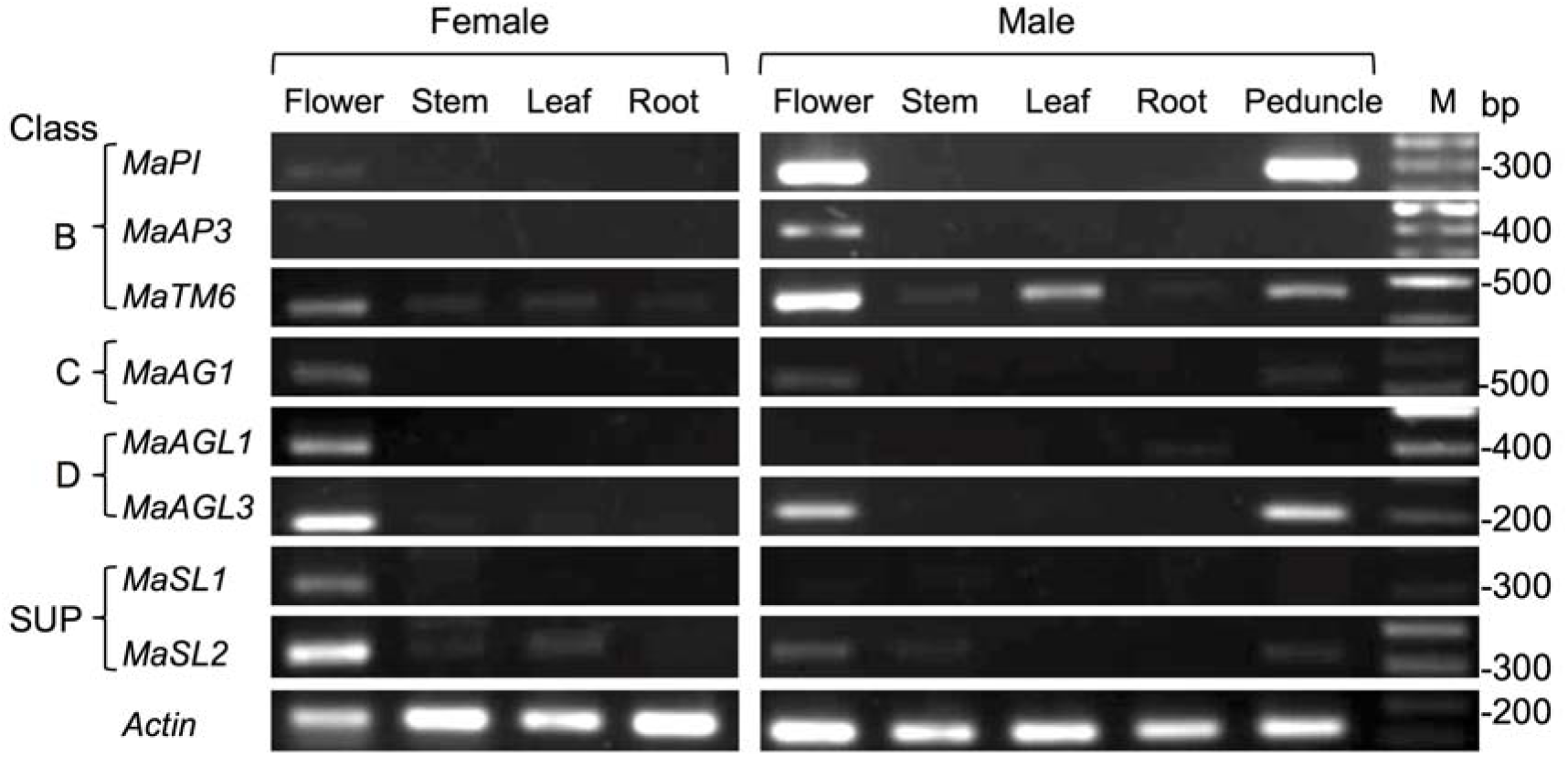
Expression pattern of floral homeotic genes in different organs of female and male plants of *M. annua*. Expression of class B, C, D and SUPERMAN-like genes was determined using semi-quantitative PCR using cDNAs derived from RNA prepared from the indicated organs. Actin was used as ubiquitously expressed reference gene. M, molecular size markers in base pairs.

BAP-induced feminization of male plants resulted in changes in expression patterns of floral genes (Fig. 5). The expression of the class B identity gene *MaTM6* was not significantly affected by feminization, while *MaPI* and *MaAP3* were down-regulated. In contrast, the expression of class C/D floral genes, namely, *MaAG1, MaAGL1* and *MaAGL3* as well as *MaSL1* was up-regulated by feminization. A significant up-regulation of class C and D genes was observed at 8-11 days of BAP treatment.

**Figure 5.**
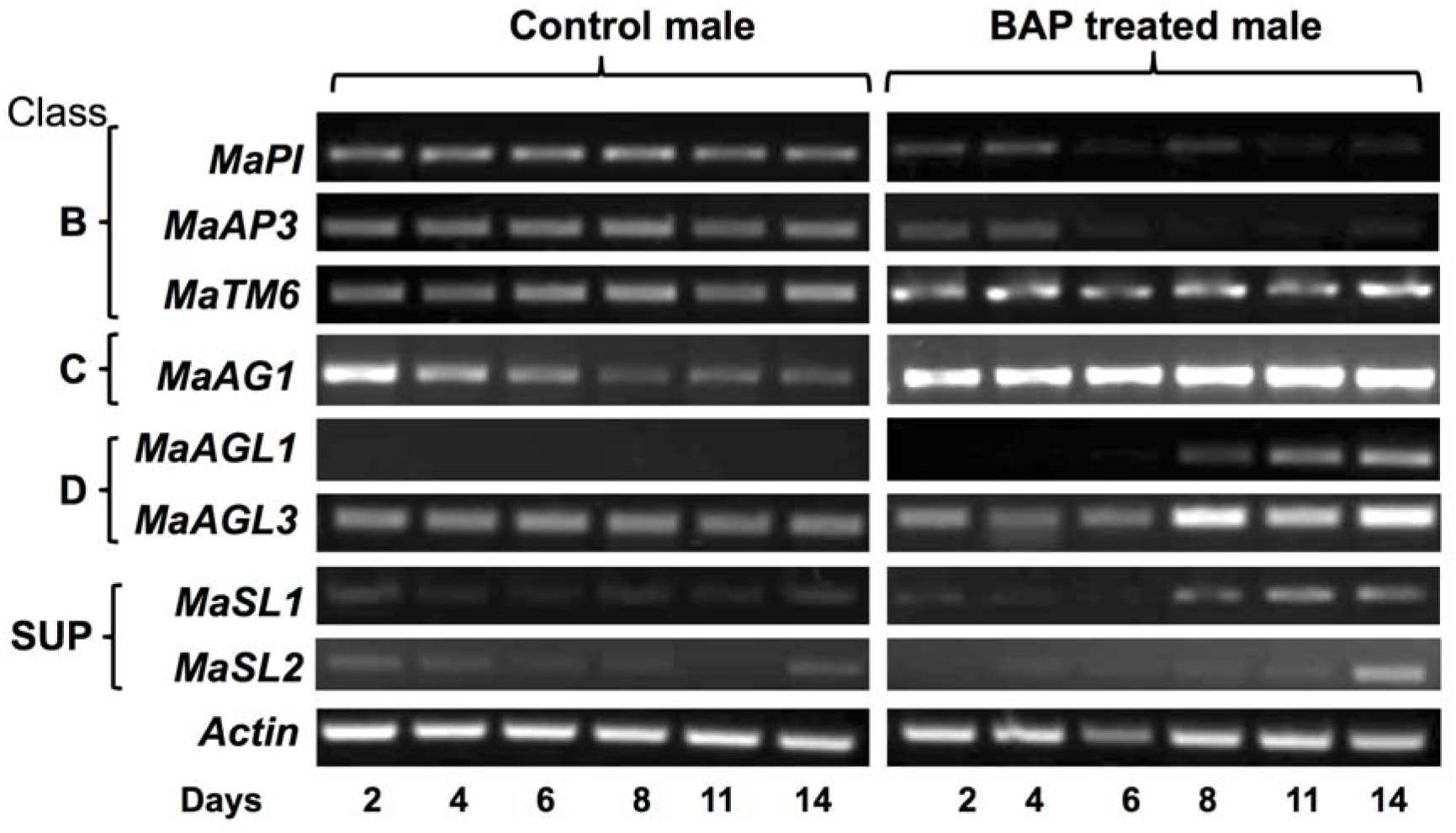
A time course of the expression of floral genes during BAP-induced feminization. 25-day-old *M. annua* plants were sprayed 3 times daily with water (control male) or with cytokinin (BAP-treated male). Newly emerging inflorescences were collected on the indicated days, RNA was prepared and subjected to cDNA synthesis. The expression of the indicated floral genes was determined by semi-quantitative PCR using cDNAs as templates. The class to which floral homeotic genes belong is indicated on the left. Actin was used as reference gene.

### Epigenetic regulation of floral genes

Epigenetics has often been implicated in sex determination in dioecious plants (Janousek *et al*., 1996; Brautigam *et al*., 2017). We thus wanted to address the involvement of epigenetic mechanisms in the regulation of floral gene expression. To this end, we first examined the chromatin configuration of promoters of several floral genes by micrococcal nuclease (MNase) assay. The MNase-treated nuclei from male and feminized male flowers (after 14 days of BAP treatment) showed similar progressive digestion of genomic DNA with notable nucleosomal ladders (Fig. 6A). MNase-digested DNAs was used as templates for PCR analysis of promoter regions of several genes. The results showed (Fig. 6B) two major digestion pattern reflecting open and relatively close chromatin configuration. Yet no notable differences in digestion pattern could be observed between male and feminized male flowers. Accordingly, group I consists the promoter regions of class B genes *MaPI* and *MaAP3* showing higher sensitivity to MNase digestion similarly to actin, a constitutively expressed gene. Group II, which composed of the class D gene *MaAGL1* as well as *MaSL1* and *MaSL2* were more resistant to MNase digestion (Fig. 6B). Thus, it appears that class B genes that assume an open chromatin conformation in male flowers remained open upon feminization, while class D assumes a relatively close configuration in male and feminized male flowers.

**Figure 6.**
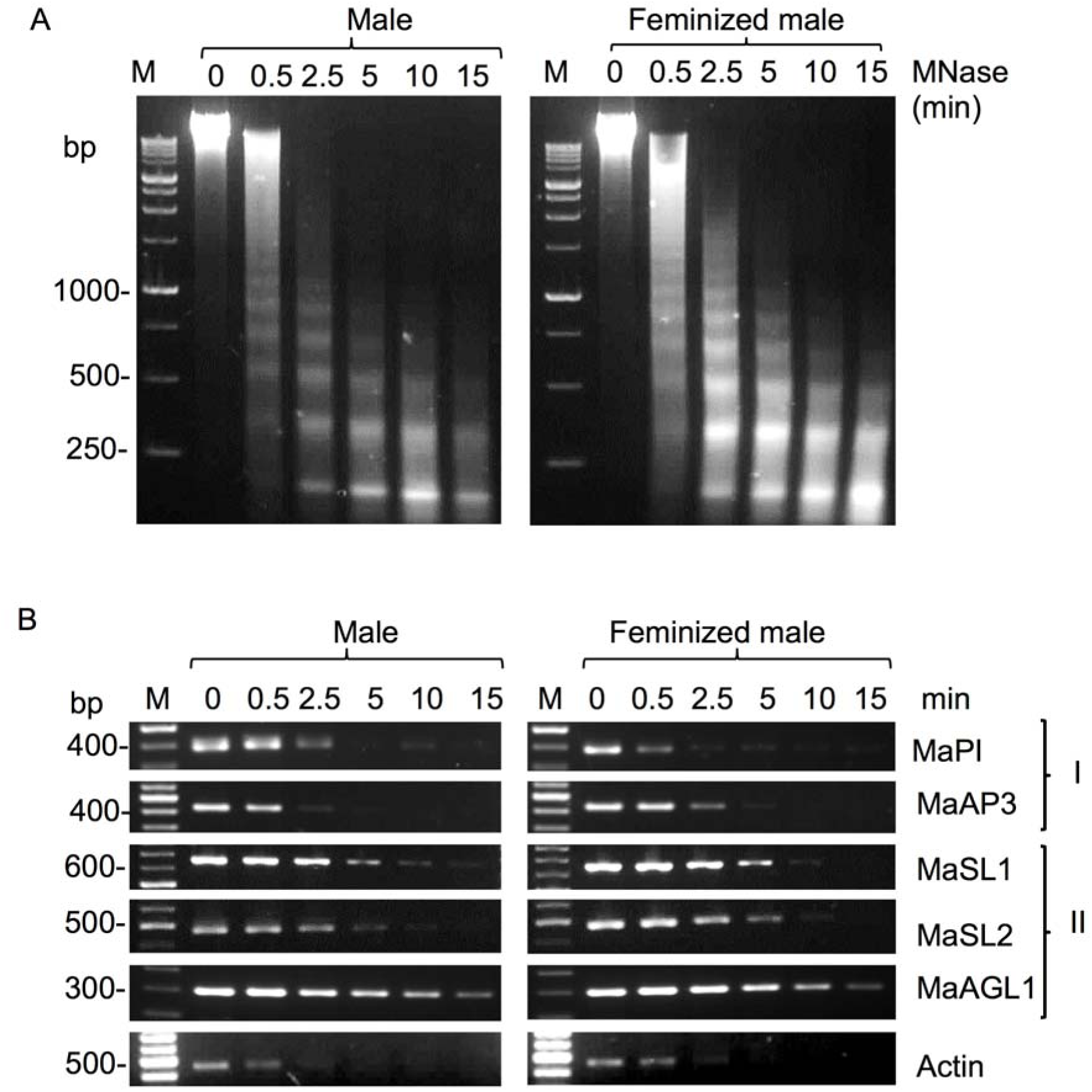
Analysis of chromatin configuration of selected floral genes by micrococcal nuclease assay. (**A**) Nuclei prepared from male and feminized male flower buds (BAP-treated for 14 days, before female flowers are visible) were treated with MNase for the indicated time periods. DNA was extracted from MNase-treated nuclei and resolved on 1.5% agarose gel. M, molecular size marker in base pairs. (**B**) Assessment of chromatin configuration of promoters of the indicated genes was performed by PCR using DNA recovered from MNase-treated nuclei (shown in A). Group I refers to male-related identity genes and Group II to Female-related identity genes. Actin was used as reference for open chromatin configuration. M, molecular size markers in base pairs.

To examine the role of DNA methylation in the control of chromatin configuration and expression of floral genes, the status of cytosine methylation at the promoter regions of several differentially expressed genes, namely, *MaSL1, MaSL2* and *MaAGL1* was assayed by chop-PCR using the methylation sensitive enzymes *Hpa*II and *Msp*I. Notably, both enzymes recognize the CCGG site but differ in their sensitivity to cytosine methylation. While *Hpa*II is sensitive when either of cytosine is methylated, *Msp*I is sensitive only when the external cytosine is methylated, allowing distinguishing between CG and CHG methylation. Chop-PCR revealed no differences in CpG methylation status of the examined genes in female and male flowers. However, CHG methylation appeared to be absent from the promoter regions of *MaAGL1* and *MaSL2* genes in male flowers inasmuch as no recovery of PCR fragment could be detected from *Msp*I digest (Fig. 7A). We also perform bisulfite sequencing of *MaAP3* and *MaSL1* promoter and gene body regions showing no differences in DNA methylation status between male and female flowers. The promoter regions of both genes were highly methylated at all cytosine contexts (CG, CHG and CHH) while their gene bodies were essentially unmethylated (Fig. 7B).

**Figure 7.**
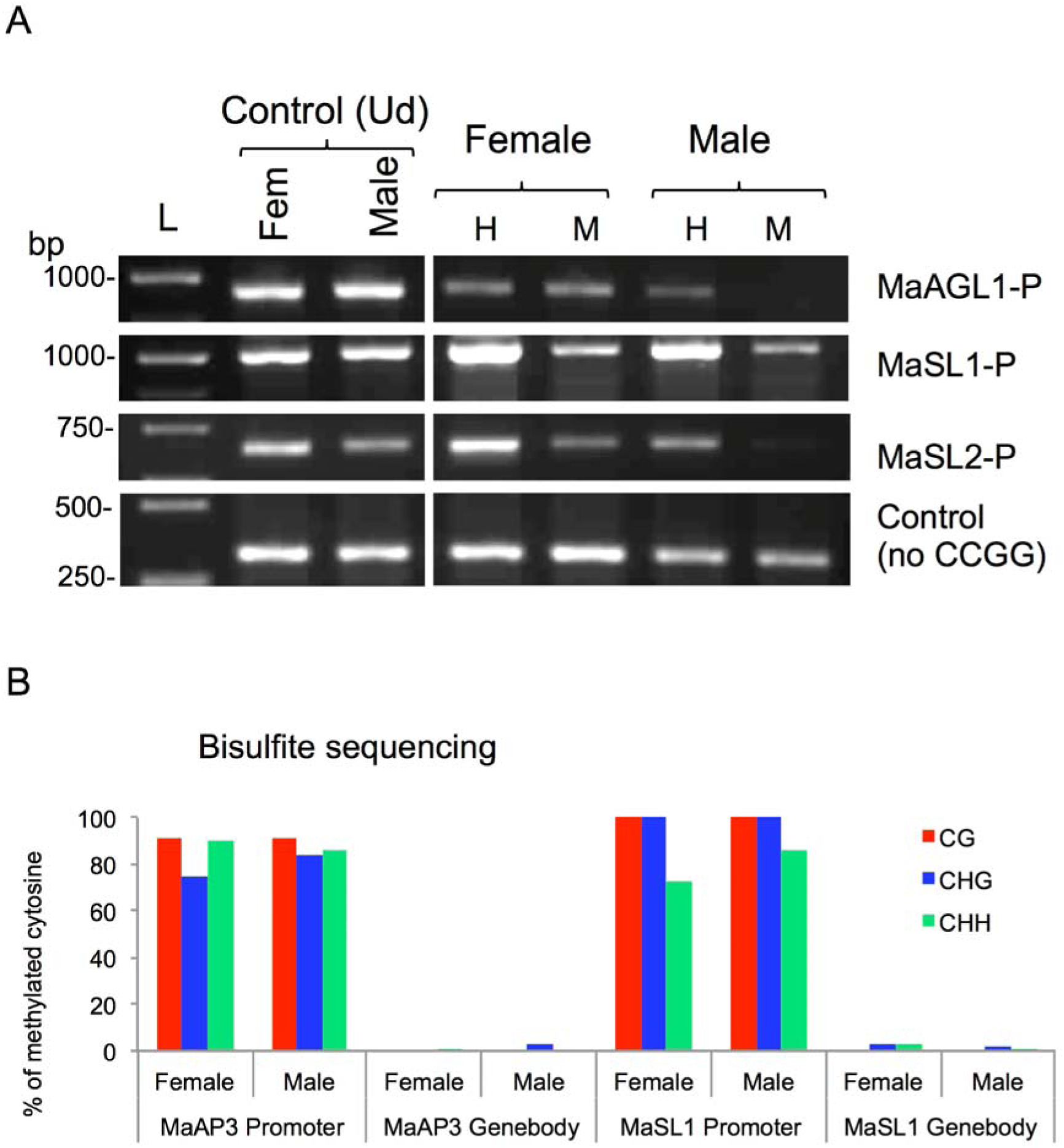
Transcriptionally active floral genes are methylated in both genders of *M. annua.* (**A**) Analysis of DNA methylation at promoters of MaAGL1, MaSL1 and MaSL2 genes by Chop-PCR. A fragmet of the *MaSL1* gene lacking CCGG site was used as control. Left panel is a control of undigested DNA (Ud). H, *HpaII*; M, *MspI*; L, molecular size markers in base pairs. (**B**) Analysis of methylation at promoter and gene-body of *MaAP3* and *MaSL1* genes by bisulfite sequencing. The percentage of cytosine methylation for each fragment was determined from at least 10 different clones.

## Discussion

The annual dioecious *Mercurialis annua* is a unique experimental system to study mechanisms underlying plant sexual dimorphism. A major advantage of this species is the possibility to feminize male plants that produce viable seeds. The change of the fate of the male flower development by cytokinin treatment of the plants (Louis *et al*., 1990; Duran and Durand, 1991), enabled investigation of gene regulation at various levels: proteome, mRNA and epigenetics. The data obtained in this study regarding the expression of floral identity genes are consistent with their known function in determining sexual identity of floral organs in various plant species. It has been shown previously that class B genes were highly expressed in male flowers, of the type-II dioecious plants *Thalictrum dioicum* and *Spinacia oleracea* (Di Stilio *et al.*, 2005; Pfent *et al.*, 2005). In agreement, our results showed that male flowers are characterized by a strong expression of class B genes, *MaPI* and *MaAP3*, concomitantly with suppression of female identity genes such as *MaAGL1* (class D) and *MaSL1*. The involvement of cytokinin in sex determination has been reported in a variety of plant species including the oilseed crops *Plukenetia volubilis* and *Jatropha curcas* (Pan and Xu, 2011; Fu *et al*., 2014).

### Expression pattern of floral genes

Proteome analysis of BAP feminized males showed differential expression of several protein families including nucleic acid binding proteins, hydrolases, ligases, transferases and transcription factors. Interestingly, floral homeotic MADS-box gene product homologs of *Arabidopsis* class E genes, SEP1, SEP3 and class D gene SHP2/AGL5 were up-regulated and homolog of class B gene, PI, involved in specification of petals and stamens, was down-regulated in feminized males. The proteins, SEP1 and SEP3 were implicated in regulation of all four flower whorls of *Arabidopsis* (Zahn *et al*., 2005); while in other plants species, *SEP*-like genes play diverse roles in growth and development including plant architecture, ovule development, fruit ripening, inflorescence architecture and reproductive meristem fate (Uimari *et al*. 2004; Biewers, 2014). In *Gerbera hybrida* two duplicated orthologs of SEP-like gene *GRCD1* and *GRCD2* were sub-functionalized for stamen and carpel identity, respectively. The *Mercurialis* orthologs of SEP1 and SEP3 proteins presented here might have a role in female flower specification. The up-regulation of SHP2/AGL5 in feminized males is consistent with SHP role in carpel development in *Arabidopsis*. Accordingly, constitutive expression of SHP genes in *Arabidopsis* resulted in a partial conversion of the first whorl sepals into carpel-like structures demonstrated by extensive proliferation of stigmatic papillae (Favaro *et al*., 2003; Pinyopich *et al*., 2003). The PI protein, which was down-regulated in feminized males was involved in controlling the development of whorls 2 and 3 in *Arabidopsis*, *Antirhinum* and tomato (Trobner *et al*., 1992; Goto and Mayerowitz, 1994; Guo *et al*., 2016). Thus, these results suggest that the cytokinin switched-off the male control genes (e.g., *PISTILLATA*) concomitantly with up-regulation of female identity genes consequently leading to the replacement of stamen by carpels, as in the development of normal dioecious female flower.

An earlier study, using a cell-free translation system with RNAs derived from *M. annua* male and female flowers demonstrated peptide variation between the two sexes and that cytokinin-induced feminization of male flowers has led to the expression of female-specific peptides (Deligue *et al*., 1984). Similarly, we found that cytokinin-induced feminization of *M. annua* male flowers was associated with upregulation of female-specific floral genes concomitantly with downregulation of male-specific genes. The effect of cytokinin on floral gene expression was reported previously (Estruch *et al*., 1993). Accordingly, the expression of the cytokinin-synthesizing gene *IPT* in transgenic tobacco plants resulted in abnormal flower development concomitantly with a notable decrease in accumulation of class B genes (*DEFA*, *GLO*) and class C gene (*PLENA*) (Estruch *et al*., 1993). In *Arabidopsis*, exogenous application of BAP was reported to promote differentiation of carpeloid tissue and suppress stamen development. This is similar to the effect obtained by overexpressing *SUP* in tobacco plants leading to the proposition that *SUP* may regulate sex determination pathways by promoting female organ differentiation *via* its effect on cytokinin signaling (Nibau *et al*., 2011). Alternatively, cytokinin may affect male and female flower development *via* controlling *SUP* expression. Indeed, in *M. annua* as well as in the dioecious *Popolus tomentosa* and *Silene latifolia* the *SUP*-like genes exhibited female flower-specific expression (Kazama *et al*., 2009; Song *et al*., 2013). In *Arabidopsis*, *sup* mutant was associated with the ectopic expression of *AP3* gene in the fourth whorl (Bowman *et al*., 1992), therefore *SUP* was proposed to function as a negative regulator of *AP3*. The concomitant expression of class B and SUP-like genes in male flower buds suggests that *SUP*-like gene(s) might not be a transcriptional regulator of class B genes in *M. annua*. An alternative possibility is that the *SUP* gene expression in male flower buds is negatively regulated post-transcriptionally.

The expression of class B gene, *MaAP3*, was restricted to male flowers, while *MaTM6 (AP3-*related) and *MaPI*, were expressed in flowers as well as in peduncles (Fig. 4). It is noteworthy that *TM6*, which is absent in *A. thaliana*, was also expressed in leaves of *M. annua* male plants and weakly in other vegetative organs. The broader expression pattern of *TM6* orthologs was reported in *Carica papaya* (*CpTM6-2*) and *Vitis vinifera* (*VvTM6*); *CpTM6-2* was expressed at a low level in sepals and at a high level in leaves (Ackerman *et al*., 2008), while *VvTM6* was expressed throughout the plant, though displaying high levels in flowers and berries (Poupin *et al*., 2007). It has been proposed that a gene duplication event of the *paleoAP3* genes resulted in two types, *euAP3* and *TM6* lineages that are distinguished by their C-terminal regions (Kramer *et al*., 1998). These duplicated genes probably adopted, to some extent, different functions (sub-functionalization) demonstrated by their tissue-specific patterns of expression and the effect of their loss-of-function on flower development (Eckardt, 2006).

The expression of class C gene, *MaAG1* was similar in male and female flowers of *M. annua* suggesting it may not involve in gender determination. This is consistent with previous reports showing that the C class *AG* genes are involved in the floral quartet specifying both stamens and carpels (reviewed in Theissen *et al*., 2016). The class D genes, *MaAGL1* and *MaAGL3* were highly expressed in female flowers; *MaAGL3* was also expressed in male flowers as well as in peduncles (Fig. 4). Our results suggest that class B genes *MaAP3*, *MaPI* together with class C *MaAG1* have a role in determining the identity of male floral organs. These gene products may participate in the floral quartet that controls gene expression and male reproductive organ identity (Theissen *et al*., 2016). On the other hand, class D genes *MaAGL1* and *MaAGL3* together with class C and class E genes may form a floral quartet that specifies female floral organs, carpels and ovules. Notably, in seed plants the class B genes have been suggested to have a primary role in sex-determination (Winter *et al*., 1999). Accordingly, expression of both, class B and class C genes specify male reproductive organs while the expression of only class C genes specify female reproductive organs. Thus, switching from male to female and *vice versa* can be solely driven by changes in the spatio-temporal expression of class B genes (Winter *et al*., 1999; Theissen and Melzer, 2007). Our data however showed that induction of feminization was associated not only with turning off expression of class B genes but also with up-regulation of class C/D genes, which might be crucial for the development of female flower in otherwise male plants of *M. annua*.

### Epigenetics and sex determination

Our data showed that there is no clear relationship between floral homeotic genes and their epigenetic makeup (Table 1). Gene expression is primarily regulated at the chromatin level where gene transcription requires open chromatin to allow for the transcription machinery to approach the gene locus. The analysis of chromatin accessibility by MNase assay revealed that in male flowers class B genes *MaPI* and *MaAP3* assume an open chromatin conformation similar to the constitutively expressed gene *Actin*. On the other hand, class D genes *MaAGL1* and SUP-like genes *MaSL1* and *MaSL2* appeared to acquire a relatively close conformation, which is consistent with the lack of expression in male flowers. Surprisingly, however, upon feminization and up-regulation of *MaAGL1* and *MaSL1* no apparent change in accessibility of chromatin to MNase was evident. This suggests that chromatin can assume different levels of open chromatin conformation that provide another regulatory layer for control of gene expression (Ishihara *et al*., 2010; Kotomura *et al*., 2015). Similarly, no change in chromatin accessibility was observed for the down-regulated class B genes, *MaPI* and *MaAP3*, whose transcription was possibly halted in an open chromatin environment by other means (e.g., suppressor proteins).

**Table 1:** Summary of the expression level of floral genes in relation to their epigenetic constraints. -, no expression; +, low expression; ++/+++, high expression; mAll, methylated at all C context; mCG, methylated at the CG context only; S, silent; E, expressed; e, low expression; O, Open chromatin; PO, Partial open chromatin.

| Gene name | expression |  |  | DNA methylation |  |  | Sensitivity to Mnase |  |  | Score |  |  |
| --- | --- | --- | --- | --- | --- | --- | --- | --- | --- | --- | --- | --- |
|  | Female | Male | F-male | Female | Male | F-male | Female | Male | F-male | Female | Male | F-male |
| MaPI | - | ++ | - |  |  |  |  | high | high | S | O E | O S |
| MaAP3 | - | +++ | - | mALL | mALL |  |  | high | high | m S | O m E | O S |
| MaActin | +++ | +++ | +++ |  |  |  |  | high | high | E | O E | O E |
| MaAGL3 | +++ | + | +++ |  |  |  |  |  |  | E | e | E |
| MaAGL1 | +++ | - | ++ | mALL | mCG |  |  | low | low | m E | PO m S | PO E |
| MaSL1 | ++ | - | ++ | mALL | mALL |  |  | low | low | m E | PO m S | PO E |
| MaSL2 | ++ | - | - | mALL | mCG |  |  | low | low | m E | PO S | PO S |
| MaAG1 | +++ | ++ | +++ |  |  |  |  |  |  | E | E | E |

The nature of gene regulation by DNA methylation is not fully understood, but generally DNA methylation has been implicated in regulating chromatin structure and function (Niederhuth and Schmitz, 2017). DNA methylation was detected at promoters but not at gene-bodies of the examined floral genes. Interestingly, methylation status of all tested genes was similar in both sexes despite of their differential expression. In *Arabidopsis*, gene methylation was reported to correlate with gene expression level; gene-body methylation was correlated with constitutively and highly expressed genes, while promoter methylation was correlated with weakly expressed genes, which are usually tissue-specific (Zhang *et al*., 2006; Zilberman *et al*., 2007). However, in this work, a consistent correlation between DNA methylation and expression of the floral genes in *M. annua* was not found (Table 1 and Fig. 7). The floral genes *MaAP3*, *MaAGL1*, *MaSL1* and *MaSL2* were normally transcribed in spite of being heavily methylated at their promoters. Thus, it appears that DNA methylation at promoter regions of *M. annua* floral genes had a positive effect on floral gene expression, in contrast to the commonly known effect of suppression of expression by methylation, particularly when transposable elements are concerned (Lisch, 2009). This finding can plausibly be explained by lowering the affinity of repressors to their binding sites by DNA methylation. There are indeed reports that showed likewise that DNA methylation at promoters contributes to transcriptional activation of certain tissue-specific genes (Neissen *et al*., 2005; Weber *et al*., 2007; Rishi *et al*., 2010; Bahar Halpren *et al*., 2014).

We concluded that determination of sex organ identity in *M. annua* does not primarily involve epigenetic regulation of floral homeotic genes. Rather, the gender identity of a dioecious flower seems to be controlled up-stream in the regulatory pathway by a gender-specific regulator(s) that affects hormonal homeostasis. This is further supported by a recent report that identified only a handful number of epigenetically-regulated genes within the sex-determining region of *Populus balsamifera* (Brautigam *et al*., 2017). In that work it was shown that both the promoter and gene body of PbRR9 were methylated. Since this gene is a member of the two-component response regulator (type-A) gene family, which is involved in cytokinin signaling, it would be important to explore further the role of genes involved in hormonal homeostasis in sex determination in *M. annua*.

## Supplementary data

Supplementary file 1

Text: Sequence and phylogenetic analysis of floral MADS-box and *SUPERMAN-like* genes

Fig. S1. Phylogenetic analysis of class B genes from *M. annua*, *A. thaliana* and various taxonomic groups.

Fig S2: Phylogenetic analysis of *AG*-like genes from *M. annua*, *A. thaliana* and various taxonomic groups.

Fig S3: Phylogenetic analysis of *SUPERMAN-like* genes from *M. annua*, *A. thaliana* and various taxonomic groups.

Table S1: List of primers used in this study Supplementary file 2 (Excel Sheets)

S1: List of proteins identified in proteomic analysis

S2: Proteins exclusively present in female flower buds that appeared following BAP treatment

S3: Proteins exclusively present in male flower buds that disappeared following BAP treatment

## Acknowledgements

We acknowledge the support of the Albert Katz International School for Desert Studies for the graduate scholarship of JK. Also, The Blaustein Center for Scientific Cooperation (BCSC) and the PBC Program of Israeli Council for Higher Education for post-doctoral fellowships to NSY. This project was supported by the Frances and Elias Margolin Trust and by ICA in Israel, as well as by The Hans-Fischer-Gesellschaft, Munich and BAYHOST funding provided by the Bayerisches Staatsministerium für Bildung und Kultur, Wissenschaft und Kunst.

